# *De Novo* Mutations Resolve Disease Transmission Pathways in Clonal Malaria

**DOI:** 10.1101/213397

**Authors:** Seth N Redmond, Bronwyn M MacInnis, Selina Bopp, Amy K Bei, Daouda Ndiaye, Daniel L Hartl, Dyann F Wirth, Sarah K Volkman, Daniel E Neafsey

## Abstract

Detecting *de novo* mutations in viral and bacterial pathogens enables researchers to reconstruct detailed networks of disease transmission and is a key technique in genomic epidemiology. However these techniques have not yet been applied to the malaria parasite, *Plasmodium falciparum,* in which a larger genome, slower generation times, and a complex life cycle make them difficult to implement. Here we demonstrate the viability of *de novo* mutation studies in *P. falciparum* for the first time. Using a set of clinical samples and novel methods of sequencing, library preparation, and genotyping, we have genotyped low-complexity regions of the genome with a high degree of accuracy. Despite its slower evolutionary rate compared to bacterial or viral species, *de novo* mutation can be detected in *P. falciparum* across timescales of just 1-2 years and evolutionary rates in low-complexity regions of the genome can be up to twice that detected in the rest of the genome. The increased mutation rate allows the identification of separate clade expansions that cannot be found using previous genomic epidemiology approaches and could be a crucial tool for mapping residual transmission patterns in disease elimination campaigns and reintroduction scenarios.

## Introduction

The reconstruction of transmission networks by sequencing pathogen samples is a powerful technique that has become increasingly routine in the field of genomic epidemiology. By identifying infections with shared *de novo* variation or the shortest genetic distances between samples, epidemiologists are able to reconstruct detailed transmission networks in viral and bacterial pathogens such as flu (Neher and Bedford 2015), MRSA (Harris et al. 2010; Popovich et al. 2016) or Ebola (Gire et al. 2014; Park et al. 2015). Assaying *de novo* mutations has proven to be a useful strategy in settings where epidemiological data is limited or in situations where the actual transmission pathways are difficult to identify, and it is applicable to any *measurably evolving population* (as defined by Drummond et al. 2003). Using *de novo* genetic variation to track pathogen population movement has enabled the identification of disease reservoirs underlying re-emergence (Eyre et al. 2013); assessed the relative contribution of local versus imported infections in a given setting (Azarian et al. 2016); and, allowed the identification of the rate and times of transmission across national borders (Faria et al. 2017; Metsky et al. 2017).

Despite the relevance of all these scenarios to malaria transmission and the potential impact on elimination efforts, inference of transmission networks based on *de novo* variation has not been attempted. Genomics-based transmission analysis in malaria has so far been limited to lower-resolution techniques that rely upon standing variation (Nkhoma et al. 2013; Daniels et al. 2015). However these approaches require the reassortment of common alleles via sexual outcrossing and will not show discriminatory power in regions where transmission rates are too low (in which the parasite predominantly mates with itself) or in situations where malaria has been re-introduced to a region (resulting in an entirely clonal outbreak). Indeed, even within regions with ongoing outcrossing, clonal outbreaks persisting across multiple years have been found (Daniels et al. 2015); as disease-endemic countries approach elimination these situations are likely to become more common.

Several challenges prevent the direct application of transmission network reconstruction methodologies in *Plasmodium.* In addition to complications introduced by sexual outcrossing, the larger genome of *P. falciparum* and a slow mutation rate relative to prokaryotic or viral pathogens reduces our ability to accurately resolve *de novo* mutations between individuals. The most rapidly mutating loci are found within low complexity sequence, and commonly within short tandem repeats (STRs). Mutation rates for STRs can be as high as 3.77 × 10^−2^ mutations / locus / generation (Chenet et al. 2015), many orders of magnitude higher than the estimated rate for SNPs in non-STR sequence (~1.07×10^−9^ substitutions/locus/generation (Lemieux 2015)). STR mutations are themselves associated with increased mutation rates in other organisms (Lenz et al. 2014) and low-complexity sequences in *P. falciparum* show greatly increased evolutionary rates compared to high-complexity flanking sequences (Zilversmit et al. 2010).

These highly mutable, low complexity regions within the *P. falciparum* genome are difficult to access using conventional short-read sequencing approaches. Commonly used sequencing formats (100 bp paired-end reads within ~300 bp fragments) prohibit high quality alignments in repetitive sequence, particularly when the size of the repeat region approaches or exceeds the read length. Insertion/deletion (INDEL) mutations within these regions further compound the problem, as insufficient non-repetitive flanking sequence around an INDEL variant can lead to incorrect partial alignments or a complete failure to map reads (Narzisi and Schatz 2015).

By spanning long repeats, longer read lengths will greatly improve our ability to analyse INDEL and STR variation in low complexity genomic regions and may, along with associated algorithmic advances in variant calling, enable studies of *de novo* variation in eukaryotes (Weisenfeld et al. 2014). Applying such an approach to clinical samples representing clonal expansions in Senegal, we show that *de novo* mutation can be reliably called in a eukaryotic parasite at a specificity that makes it amenable for determining phylogenetic relationships. We further demonstrate that *P. falciparum* can be considered a *measurably evolving population* and is therefore a suitable subject for transmission network reconstruction. The wider use of these techniques would greatly increase the resolution of genomic epidemiology studies in malaria and could have a dramatic impact on efforts to eliminate the disease.

## Results

To increase our ability to detect rapidly arising mutations in the *P. falciparum* genome, a modified library preparation method was employed in which large volumes of DNA were prepared without a PCR step (minimizing the introduction of amplification errors); 250bp Illumina reads were generated on a size-selected 450bp fragment (ensuring overlapping reads that could be joined into one long fragment); these reads were genotyped with DISCOVAR, an algorithm designed specifically for this data type (Weisenfeld et al. 2014). The extent and accuracy of these techniques were assessed using laboratory isolates of well-defined parasite clones, each with an associated genome assembly. These techniques were then applied to patient samples collected from a single clinic in Thiès, Senegal representing 3 separate clonal expansions circulating in the region. These clonal samples were genotyped using a 24 SNP barcode (Daniels et al. 2015) and were previously determined to be identical by fixed locus genotyping (Bei *et al.* in preparation).

### 250 bp Illumina Reads Increase Genotype Specificity in Low-Complexity Regions

#### 1) Genome Accessibility

DISCOVAR and GATK HaplotypeCaller were assessed using an approach in which we generated an *in silico* library from one genome (Dd2) aligned this to another reference genome (3D7) and assessed our ability to accurately reconstruct the second reference genome. The great majority of the *P. falciparum* genome was found to be ‘accessible’ to the DISCOVAR genotyping software, apart from mitotically recombining telomere and subtelomere sequence and intercalary heterochromatin (Flueck et al. 2009). We find that 66.0% of the total genome is accessible via HaplotypeCaller using 100 bp reads as compared to 80.2% via DISCOVAR using 250 bp reads – this difference represents an additional 3.3 Mb of callable genome, predominantly in low-complexity regions where rapidly mutating loci are most likely to be found [see Fig 1]. Of 10.8 Mb of sequence identified as low-complexity by DustMasker (Morgulis et al. 2006), 7.8 Mb is accessible via DISCOVAR vs. 6.8 Mb accessible using HaplotypeCaller.

**Figure 1.**
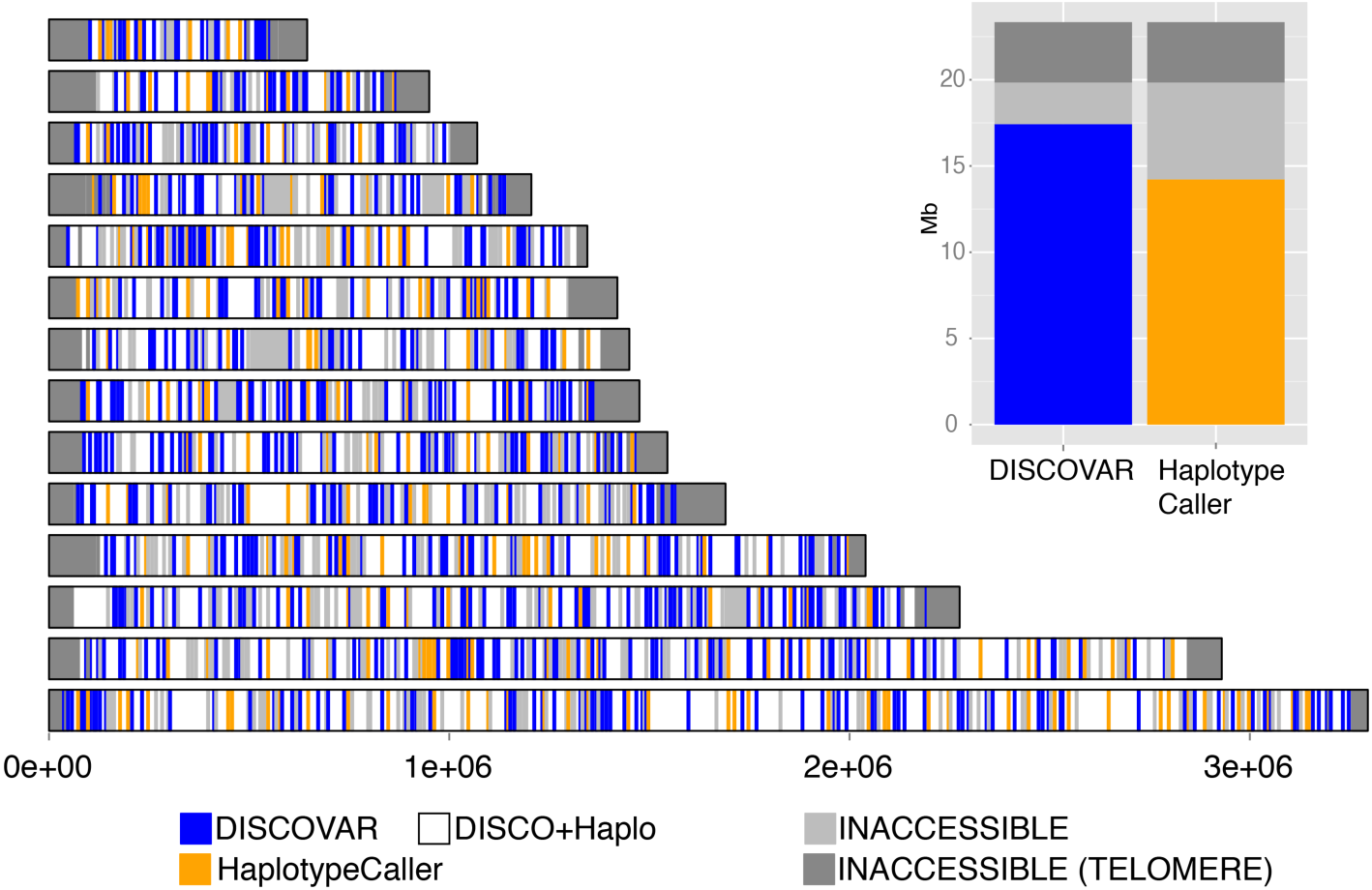
Genome accessibility was assessed based on *in silico* comparisons of two *P. falciparum* genome assemblies. Variants were called from reads simulated from the Pf_Dd2 reference and aligned to the Pf3D7_v3 reference and assessed as correct or incorrect based on a 200 bp flanking region on either side. Blocks of 1 kb across the genome were defined as accessible only if they had a read coverage within two standard deviations of the median and if they had called all variants accurately. Comparisons between GATK HaplotypeCaller and DISCOVAR indicated an increase in the proportion of the genome that was callable from 66.0% to 80.2%. Of the 10.8 Mb of low complexity sequence identified by DustMasker, 7.8 Mb was found to be accessible by DISCOVAR.

#### 2) Marker Validation

In regions that were deemed accessible to both methods, specificity and sensitivity were assessed by comparing calls to the Pf-Crosses variant set for these two strains; Receiver-Operator Characteristic (ROC) curves were generated for accessible and inaccessible regions for both SNPs and INDELs [Fig S1]. DISCOVAR demonstrated far greater specificity compared to HaplotypeCaller, albeit at a significant cost to sensitivity (DISCOVAR specificity = 0.987, sensitivity = 0.460; HaplotypeCaller specificity = 0.374, sensitivity = 0.794). These sensitivity/specificity rates were remarkably similar when this was extended to the low complexity genome (DISCOVAR specificity = 0.982, sensitivity = 0.358; HaplotypeCaller specificity = 0.344, sensitivity = 0.780). In the DISCOVAR-only regions of the genome an excess of false positive SNPs was seen when calling with DISCOVAR; in comparison the true-positive: false-positive ratio of GATK HaplotypeCaller (i.e. the ratio of variants found in the Pf-Crosses dataset to those that were not) remained constant in both the core and ‘HaplotypeCaller-only’ regions.

#### 3) Accessible / Inaccessible Genome Comparison

The higher specificity genotyping techniques were applied to a clinical isolate collection from Senegal, which were indistinguishable by previous techniques and represented independent infections collected over multiple years. Due to the lack of pre-existing dataset exhibiting high-quality genotypes in low-complexity regions that could be used to validate the DISCOVAR calls, we instead assessed genotype call accuracy by comparing genetic distances between the samples, examining the correlation between genetic distances as calculated for different variant types and regions [Fig S2]. Even if INDELs mutate at a different rate from SNPs, or low complexity regions show a higher overall mutation rate than other loci, we still would expect a high correlation between these two measures across an entire population. Genetic distances were compared between the regions of the genome accessible to both callers and those regions accessible to DISCOVAR-only; correlation was high and statistically significant (R = 0.91, *p* < 1e15 – calculated by Pearson's product moment) [Fig S3]. Similarly, correlation between SNP and INDEL distances for the *de novo* variants (within each clade) is strong, and is higher in the DISCOVAR distances (*R* = 0.95) than it is in the GATK distances (*R* = 0.89, *p* < 1e15) [Fig S4]. We conclude from this that both INDELs and SNPs in the low-complexity regions are suitable substrates for phylogenetic inference.

#### 4) Higher Specificity Genotyping Methods Allow Evolutionary Inference in Low-Complexity Regions

Variants generated with either HaplotypeCaller or DISCOVAR were used to generate a phylogenetic tree via maximum parsimony, with high bootstrap values in both cases (mean 0.71 HaplotypeCaller, 0.78 DISCOVAR) representing a similar level of support for the best-supported phylogeny using either genotyping method. Many splits were found in both phylogenies. However, even though a far greater number of parsimony-informative (N ≥ 2) markers were called using HaplotypeCaller, we conclude that the use of 250 bp reads and DISCOVAR genotyping produced superior phylogenies. Lento / bipartition plots were generated for each phylogeny showing the proportion of variants that either supported each bipartition, contradicted it (requiring a homoplasy or reversion to explain the split), or were irrelevant (e.g. a variant contained entirely within one clade). Despite similar bootstrap values for the two data types, the proportion of markers supporting the best phylogeny was higher for the DISCOVAR-derived calls (where for each split in the phylogeny, 1-4% of 1858 variants supported alternative splits) than it was for the HaplotypeCaller calls (3-32% of 8290 variants supported an alternative split). The majority of the DISCOVAR genotypes that were not irrelevant supported the major nodes in the tree (mean 74.4% support of relevant genotypes), whereas only a minority of the HaplotypeCaller genotypes did (mean 18.3% support) [Fig S6]. It should be noted that these non-supporting variants did not support any single discordant topologies, and reinforces the conclusion of our ROC analyses [Fig.S1] that the GATK HaplotypeCaller calls contained a far higher number of false positive variants. Due to the superior specificity of DISCOVAR genotyping compared to HaplotypeCaller, this method was used for all subsequent analyses.

### *De Novo* Variants Can Distinguish IBD Parasites

Of the 26,696 DISCOVAR variants that distinguished the three clades from each other, 3,200 segregated within the clades (622, 828, 1858 for clades 24 26 and 29 respectively). Some loci segregated within more than one clade (9 loci in clades 24 and 26; 39 in 24 & 29; 57 in 26 & 29; 1 locus segregated in all three clades) [Fig S2]; though many of these loci showed different alleles in different clades, we are unable to distinguish between independent mutations *versus* shared variation. Genes that are frequently used for study of parasite diversity were also examined; no variants of any type were found within any of the clades for *MSP1, MSP2,* SERA2, *GLURP, TRAP* or *CSP.* The uniformity of these markers within an otherwise panmictic population indicated that each clade descended from a single common ancestor without recombination, the clades were therefore considered identical by descent (IBD). Within clades, pairwise distances ranged from 88 to 677 variants, with both extremes being present in clade 29. As might be expected, the oldest sample (Th086.07, collected in 2007) exhibited a greater genetic distance with respect to the other samples in clade 29, though this pattern was not repeated in the other two clades consisting of more contemporaneous samples (2012-2013; [Fig 2]).

**Figure 2.**
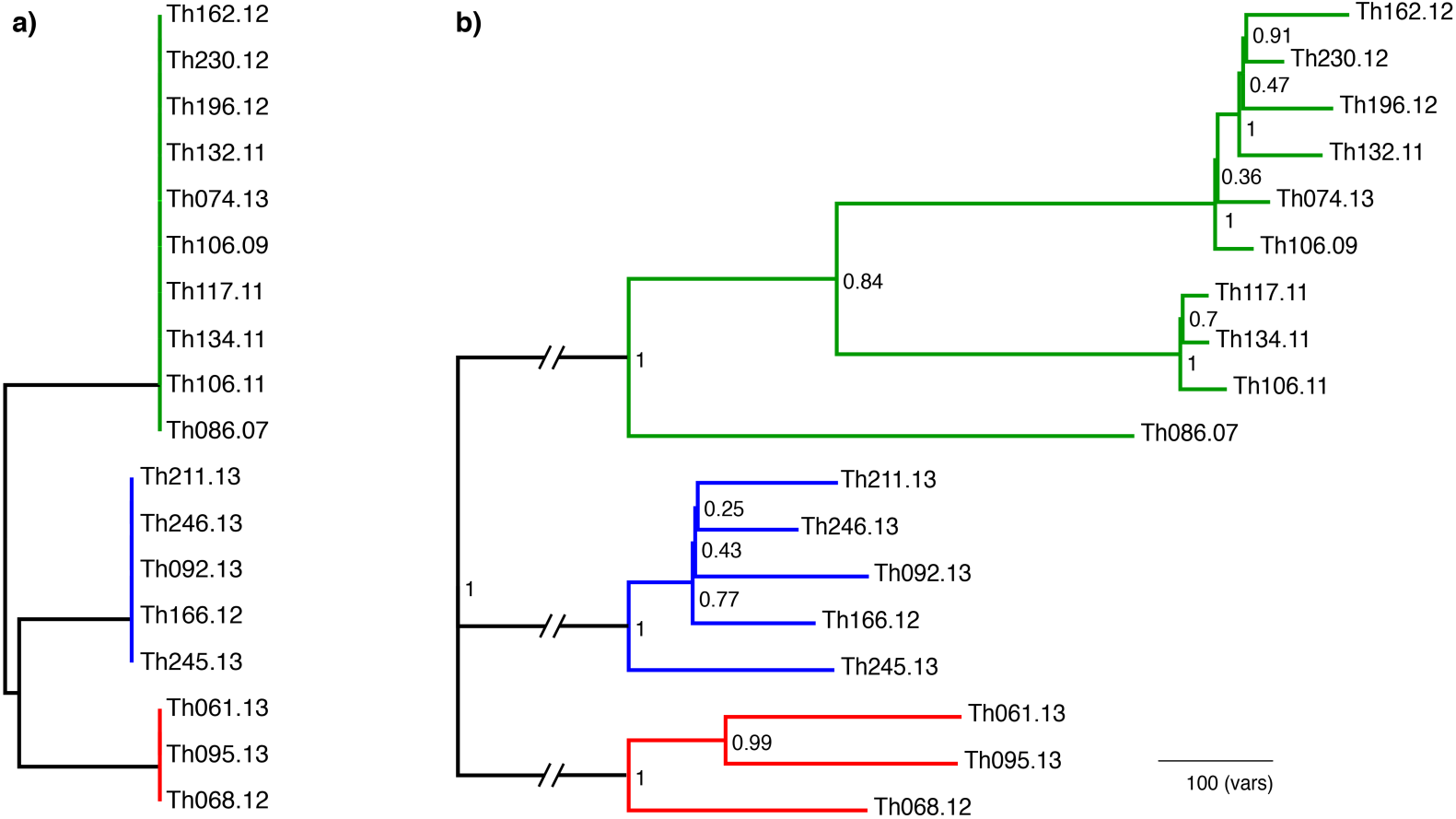
Phylogenetic resolution of parasite samples using standing variation vs. *de novo* variants. **a)** Maximum parsimony tree of standing variation (24-SNP molecular barcode - Daniels et al. 2015). The last two numerals in sample names indicate the year of collection. Clades are resolved, but samples within clades are indistinguishable. **b)** More than 3000 *de novo* variants were called via DISCOVAR that segregated the individuals into three IBD clades. Samples within clades 24 (red), 26 (blue) and 29 (green) were separated by 622, 828 and 1858, variants respectively, with the closest individuals (Th106.09 / Th074.13) distinguished by 88 *de novo* variants. Phylogenetic trees were calculated using 15276 SNPs and 11420 INDELs using maximum parsimony. Numbers on nodes indicate bootstrap support. Our ability to discern phylogeny using only *de novo* variants was high with bootstrap values above 0.5 for all nodes subtending samples collected at more than one year apart.

Variants were found predominantly in intergenic sequence for both SNPs (78.5%) and INDELs (83.7%), with in-frame INDELs and missense SNPs generating a further 15% of the variants in each class [Table 1] suggesting that selective pressure was acting on these mutations and that selection would not significantly confound phylogeny reconstruction. Perhaps surprisingly, while we would expect the majority of sequencing errors to be observed as singletons, the proportions of nonsense variants did not increase when considering only singletons, suggesting that some nonsense variants may be authentic. Indeed the proportion of intergenic variants significantly increased when considering only singletons, possibly representing large numbers of terminal-branch variants in the rapidly mutating intergenic regions.

**Table 1:** Mutation Consequences By Variant Class And Allele Frequency

| Variant Consequence | INDEL |  | SNP |  | N≥2 |  | Singleton |  |
| --- | --- | --- | --- | --- | --- | --- | --- | --- |
| frameshift_variant | 12 | (1.0%) | 8 | (0.3%) | 4 | (0.2%) | 0 | (0.0%) |
| inframe_deletion | 87 | (7.1%) | 60 | (2.4%) | 27 | (1.6%) | 0 | (0.0%) |
| inframe_insertion | 85 | (6.9%) | 76 | (3.0%) | 9 | (0.5%) | 0 | (0.0%) |
| intergenic | 1025 | (83.7%) | 1941 | (76.1%) | 1415 | (86.0%) | 2331 | (78.5%) |
| intragenic_variant | 0 | (0.0%) | 1 | (0.0%) | 0 | (0.0%) | 1 | (0.0%) |
| missense_variant | 12 | (1.0%) | 315 | (12.4%) | 142 | (8.6%) | 445 | (15.0%) |
| non_coding_exon_variant | 4 | (0.3%) | 11 | (0.4%) | 3 | (0.2%) | 10 | (0.3%) |
| stop_gained | 0 | (0.0%) | 9 | (0.4%) | 3 | (0.2%) | 12 | (0.4%) |
| synonymous_variant | 0 | (0.0%) | 128 | (5.0%) | 43 | (2.6%) | 171 | (5.8%) |

### SNP Errors Are Apparent Between Closely Related Samples

Mean SNP:INDEL ratios between clades were 2:1 as expected from previous work, however the most closely related pairs within each clade showed a markedly elevated SNP count, with SNP:INDEL ratios as high as 6:1 within clade 29 (samples Th132.11 / Th162.12 are separated by 141 SNPs and 23 INDELs) [Fig S2 / S5]. Skewed SNP:INDEL ratios were observed not only among those samples with the smallest genetic distance. For instance, the oldest sample (Th086.07) showed elevated pairwise SNP:INDEL ratios when compared to 6 of the other clade 29 samples. Significance of the SNP:INDEL skew was tested by the Fisher exact test; 25 of the pairs were significantly (*p* < 0.01) elevated as compared to the background 2:1 ratio. Transition:transversion (Ts:Tv) ratios were also elevated within these closely related pairs. Background Ts:Tv (calculated between clades) was 1.07-1.1:1, however within clades this rose to 4.38:1 for the closest pairs. Elevations in Ts:Tv ratios closely followed patterns of elevation of SNP:INDEL ratios, with both ratios being elevated in pairs with the shortest genetic distances. The elevation in Ts:Tv ratios and the lower specificity of DISCOVAR-called SNPs within the low-complexity genome leads us to conclude that the increase in SNP:INDEL ratio represents sequencing error rather than an effect of purifying selection. These errors are most apparent within IBD clades because of the low number of legitimate *de novo* mutations distinguishing members of those clades.

### Phylogeny & Within-Clade Structure

Phylogenies were calculated by maximum parsimony [Fig 1] and neighbor joining for SNPs and INDELs combined (because the SNP/INDEL evolutionary rates are not known *a priori,* Manhattan distance was used as the combined measure). Despite the relatively short sampling times covered (only six years in the largest clade) bootstrap support is above 50% for most within-clade relationships (mean bootstrap value 0.71). Those samples with low bootstrap support were separated by no more than a year. An unbalanced phylogeny might result from either super-spreader events or long infection chains (Colijn and Gardy 2014) both of which will generate more internal nodes that are connected to a single leaf descendent. Both Sackin and Colless indices were used to assess the phylogenies and both showed some elevation above expectations for a balanced tree, however none of the clade phylogenies were found to be significantly unbalanced (Blum and François 2005), nor was the overall phylogeny of all three clades, and we cannot conclude that any of these phylogenies represents a super-spreader event or a single infection chain.

While all of the samples within a clade are descended from a single common ancestor over the time since this recent common ancestor we can expect some structure to have emerged within these clades. The parsimony tree [Fig 2b] for the largest clade 29 showed two distinct sub-clades, one consisting of 3 samples (subclade 29.1: Th117.11, Th134.11, Th106.11) and another larger clade with six samples (subclade 29.2: Th162.12, Th230.12, Th196.12, Th132.11, Th074.13, Th106.09), with further subdivisions in the second clade. We conclude from this that the two clades represent distinct clonal expansions that have persisted separately within the population for at least four years, despite being transmitted within the same geographical area [Fig S9].

### Clade 29 is a Measurably Evolving Population

It is important to establish whether or not any of these clades represent a measurably evolving population (MEP): a population in which a set of samples can be seen to have measurably evolved in the time between the first and last sample. In an MEP much or all of the variation will have accrued *de novo* between the first and last sampling years rather than due to divergence from a common ancestor before sampling began; as a result, the root-to-tip distance of the resultant phylogeny should correlate with the sampling time, because samples collected later have had more opportunity to diverge from the most recent common ancestor (MRCA) (Drummond et al. 2003).

The two smaller clades were not sampled over sufficient time for us to confirm that we had detected an MEP, however clade 29 showed a weak but significant correlation between root-to-tip distance and time (*R* = 0.65, *p* = 0.04) via TempEST analysis [Fig 3] and was confirmed as being a MEP. Unlike the other two clades, clade 29 was sampled over a far greater period of time, consisting of samples from 2007 to 2013, and showed a much stronger phylogenetic signal, with TreePuzzle analysis resolving 71.4% of all quartets [Fig S10]. This clade is therefore suitable for inference of evolutionary rates and can be used to reconstruct a transmission network.

**Figure 3.**
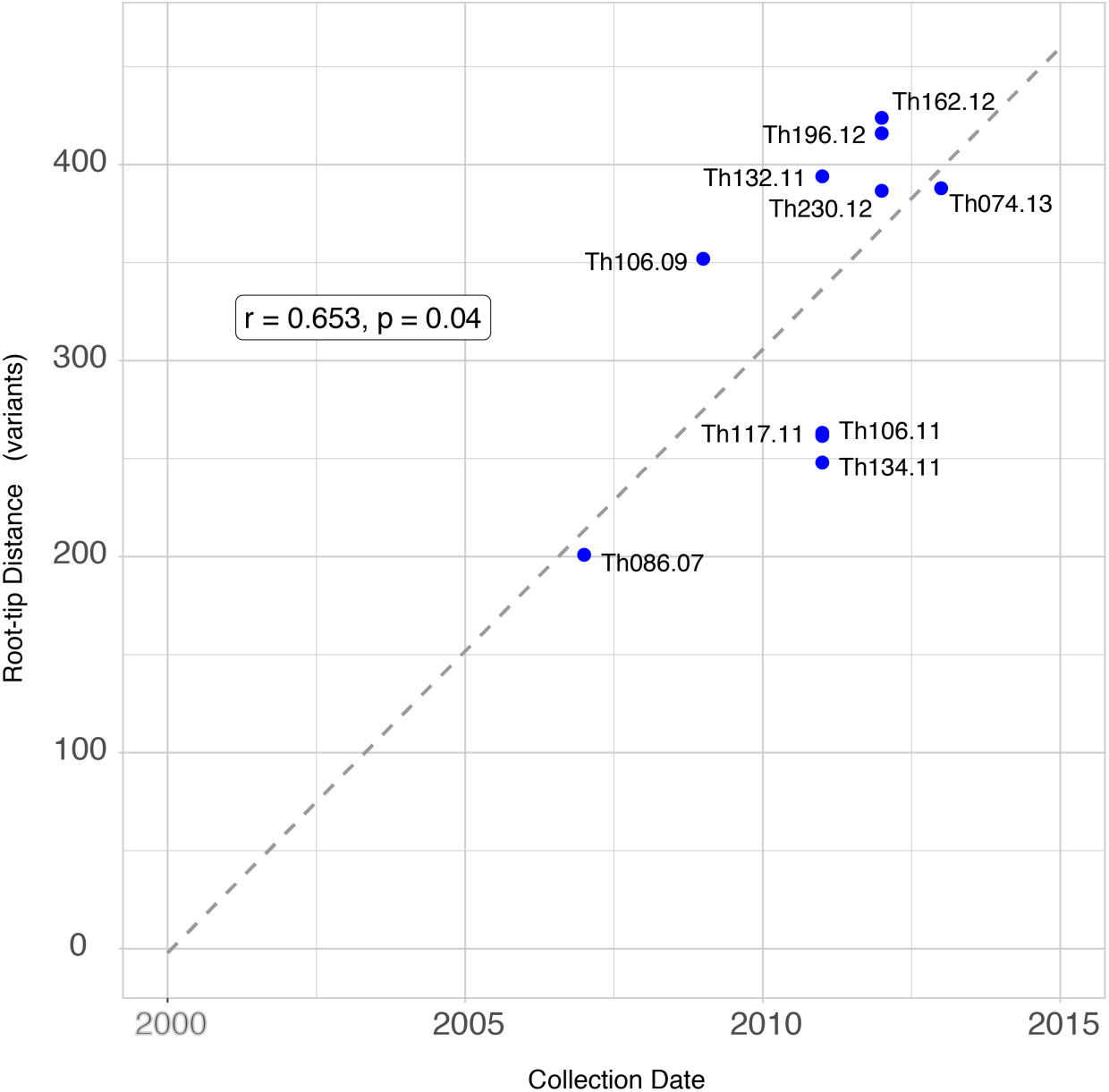
Root-to-tip distance in the phylogeny of clade 29 correlates with sampling time. The observed correlation of genetic distance and time (via Pearson’s product moment) indicates that many of our variants are *de novo* and that mutation occurs at a sufficiently high rate to resolve patterns of malaria transmission. Regression from sampling times indicates a common ancestor for the clade may have existed in approximately the year 2000.

### *De Novo* Mutations Can Reconstruct Transmission Pathways

Transmission networks were calculated using a minimum-distance based method (Jombart et al. 2011) that relied on Manhattan distances of both variant types combined [Fig 4]. Bootstrap values were high for most relationships with a majority of secondary transmission paths weakly supported. The distance-derived transmission path provided additional support for the multi-clade structure that was seen in the maximum-parsimony tree results, indicating a separate expansion of subclade 29.1 and 29.2. Both of these clades were found to be related to the basal sample Th086.07.

**Figure 4.**
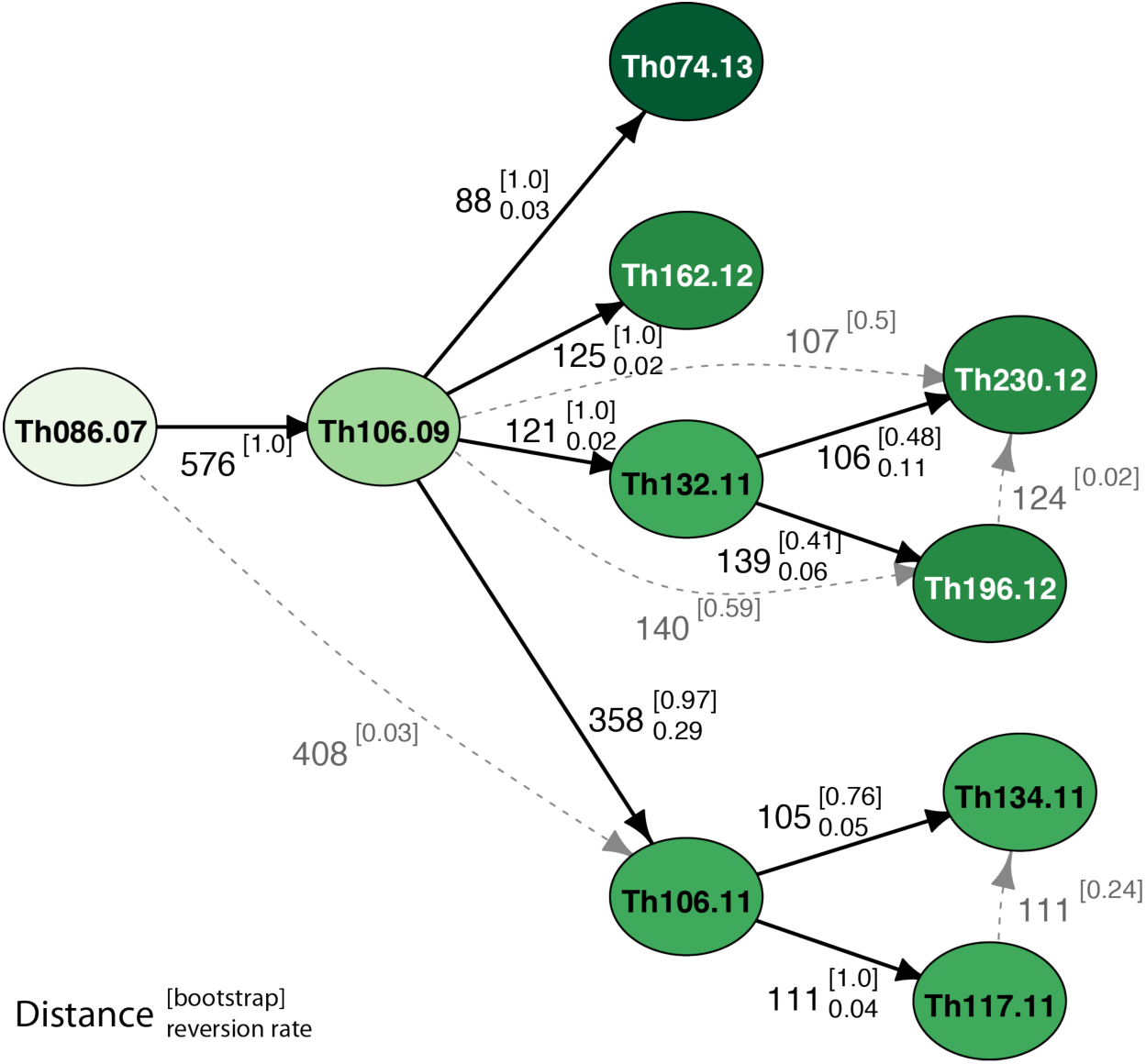
Transmission networks were estimated for clade 29 based on SNP and INDEL distances combined using a minimum distance tree approach. Edges are labeled with pairwise genetic distance, as well as bootstrap values (superscript in parentheses) and reversion rate (subscript). Bootstrap values were calculated for all edges by sampling with replacement for 100 iterations and indicate strong support for many of these distance-derived relationships. Potential alternative edges derived from the bootstrap results are shown in gray. Nodes in later years are shown in darker colours. Concordance between the parsimony tree and distance based methods is notable; in both the phylogeny and transmission network sample Th106.09 is basal to subclade 29.1 and Th106.11 to subclade 29.2. Reversion rates are significantly higher for the edge joining Th106.09 to Th106.11 supporting the independence of subclade 29.2

As a secondary confirmation of the network, we mapped individual variants to each vertex in the network and calculated the reversion rate across all primary edges (i.e., the number of *de novo* mutations that would need to be lost to support each step); we expect both variation accrued on each lineage since divergence and genotyping errors to contribute to this statistic. The majority of edges showed reversion rates of 11% or less, however the edge connecting subclade 29.2 to the rest of the samples (Th106.09 through Th106.11) showed a markedly higher reversion rate (0.29). This discrepancy indicates a lack of sampling depth in the early years of the expansion and shows that we have not captured a true progenitor of subclade 29.2.

### Evolutionary rates indicate that *de novo* genomic epidemiology studies are viable in *P. falciparum*

Evolutionary rates were calculated for all pairs within the transmission network for clade 29 using the method of Li *et al.* (Li et al. 1988). Highly variable evolutionary rates were seen between samples. Mean mutation rates were found to be 2.14 (+/− 1.6) SNPs/month/genome and 0.78 (+/− 1.3) INDELs/mth/gen [Fig S11b]. Despite the small sample set and even smaller number of IBD pairs that can be used for this analysis, it is apparent that the evolutionary rates for SNPs and INDELs are closer to the 2:1 ratio that is seen between clades than to the elevated SNP ratios seen within some of the closest samples. This strongly supports our earlier conclusion that the observed elevated SNP:INDEL ratios between closely related samples were due to sequencing errors.

Comparisons of evolutionary rate within callable and DISCOVAR-only regions of the genome show a remarkable increase in our ability to discern an MEP; the callable genome acquires 0.84 (+/− 1.8) mutations/mth/gen while the DISCOVAR-only regions gain *de novo* variants at three times this rate 2.08 (+/− 1.3) muts/mth/gen [fig S10c].

The combined evolutionary rate for all mutation classes is therefore 2.92 (+/− 2.3) muts/mth/gen [Fig 5, Fig S11a]. This remarkably high figure is comparable to the rate of mutation acquisition in some RNA viruses [Fig S12] and would indicate that most direct transmissions would have acquired at least some *de novo* mutations between infections.

**Figure 5.**
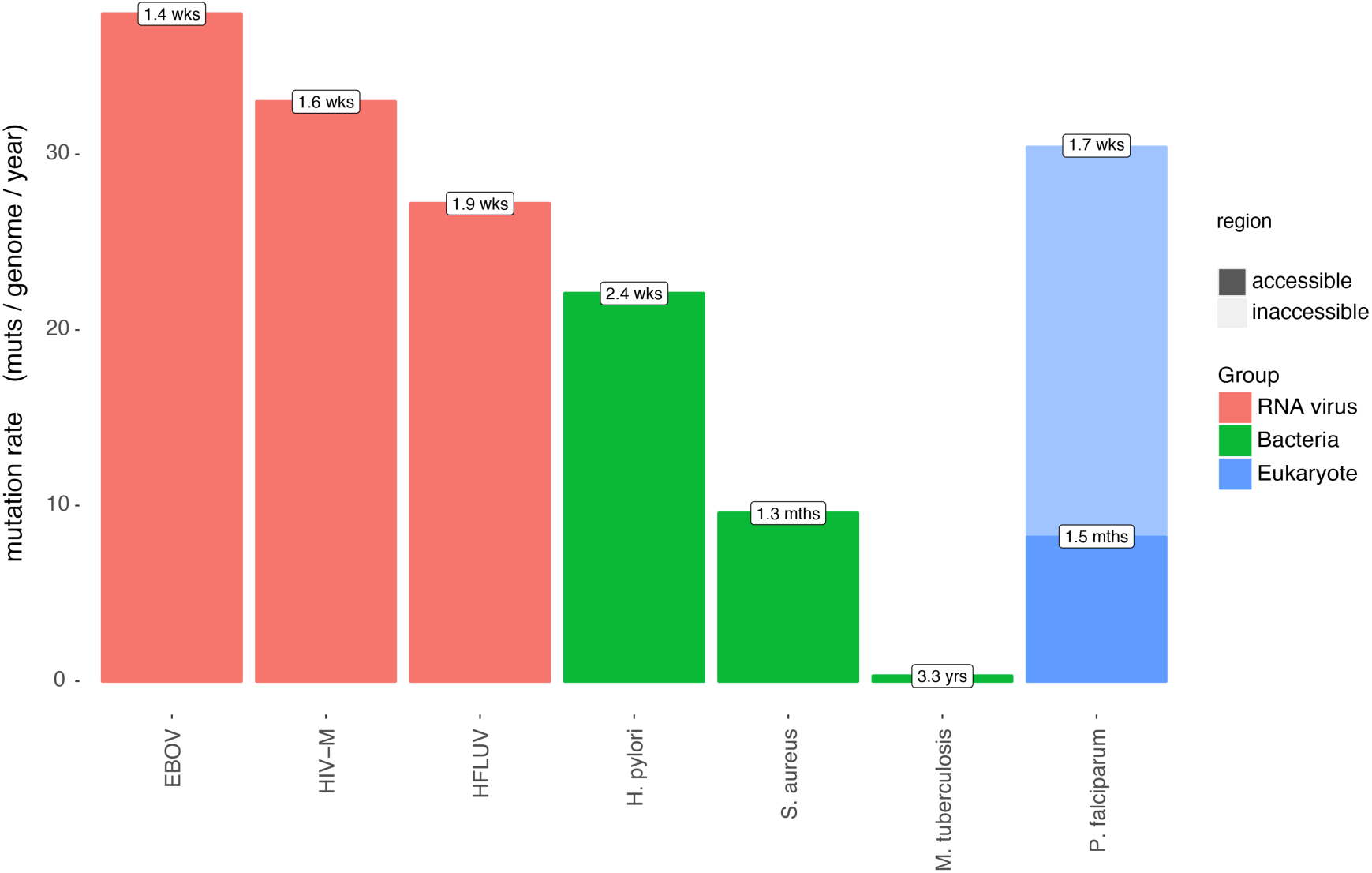
Lower mutation rates and the restricted genome size of the core genome would generate a single new mutation on average every 1.5 months. The increase in accessibility offered by DISCOVAR increases both genome size and measurable evolutionary rate, resulting in a new variant every 2.1 weeks in the low complexity genome and every 1.7 weeks overall. This makes *P. falciparum* comparable to other infectious agents like influenza or HIV where transmission networks may be informed by genome sequencing. Mutation rates derived from Biek et al. (2015)

## Discussion

This study demonstrates for the first time the viability of using *de novo* mutation to characterize transmission in a eukaryotic pathogen and further shows how this technique could be used to study the transmission of malaria—a disease that, despite a worldwide elimination campaign, still kills more than 400,000 people per year (WHO 2016). To access rapidly evolving low-complexity sequence we have used long sequencing reads, PCR-free library preparation and a large *k*-mer during the assembly stage, opening up a further 14% of the *P. falciparum* genome to accurate genotyping. The increased coverage in turn enables more rapidly mutating variants to be called than was previously possible, and the accompanying increased specificity prevents the *de novo* signal from being lost amidst the noise of sequencing or alignment error. Within a set of closely related samples from patients in Senegal (Daniels et al. 2015), we were able to call 3200 variants that successfully resolved ancestry among these parasites and generated robust phylogenies for what appeared to be identical clones when assayed at lower genetic resolution.

Two of the clades we studied were sampled across only two years each and would not have the capacity to show the correlation between sampling time and root-to-tip distance that would suggest these mutations had arisen during the sampling period. On the other hand, the third clade persisted for a total of seven years and was determined to be a measurably evolving population. This level of resolution is a first for a eukaryotic pathogen over time scales relevant to disease transmission (Biek et al. 2015) and supports the use of *de novo* variants to examine the phylodynamics of malaria in the future.

To demonstrate the utility of this approach, a transmission network was inferred for the largest clade using a minimum-distance approach [Fig 4] (Jombart et al. 2011). For characterizing malaria transmission, *de novo* variation represents a significant advance over standing variation, which would be unable to identify directional relationships between infections.

Evolutionary rates as calculated across the third-clade transmission network imply one new mutation is acquired every 1.5 weeks [Fig 5], meaning that at least one new mutation should be found between direct transmissions. While detecting a single mutation between individuals without any genotyping errors in a 23 Mb genome might be challenging with current sequencing technologies, this does indicate that sufficient *de novo* mutations to reconstruct transmission pathways can be expected in genomic epidemiology studies that were conducted over 2 years or more – times scales that are highly relevant to malaria transmission.

While the transmission network was generated as a ‘proof-of-principle’, low rates of reversion indicate that, even if the number of intervening generations remains unresolved, many of the relationships identified are accurate and some firm conclusions can be drawn from the network about the nature of transmission within Senegal.

We can eliminate the possibility that these clonal parasites persist through a single chain of infections: an infection chain would generate a tree with each node joined only to its descendent (Colijn and Gardy 2014), yet both the tree and transmission network show strong internal structure with well supported distinct clades; in particular, the clade of three samples in 2011 (Th106.11, Th117.11, Th134.11) is not directly related to the other clade found from 2011-2012 (Th132.11, Th196.12, Th230.12). It is important to note that this conclusion would have been impossible to draw from a combination of standing variation and epidemiological data: although all six of these samples were found within the Thiès region, neither clade separates geographically and individual members from separate clades were found in close proximity [Fig S9].

Perhaps more remarkably, despite 70% of the samples analyzed here having been taken in the intervening years, sample Th074.13 derives from Th106.09, and is entirely independent of the other two clades. This means that at the beginning of the dry season in 2011 (when transmission would be expected to tail off) at least three separate clades of this particular strain were circulating in the Thiès region and all survived to subsequent wet seasons. Maintenance of these parasite types suggests a sizable asymptomatic reservoir of infection may have been present.

Intriguingly, the smallest of those clades consists of just one sample Th074.13; this sample came from a village 16km outside of Thiès and is separated from its closest node in the network by six years (Table S2), yet it is separated from this parasite by just 88 variants – the shortest genetic distance in our dataset found over the longest intervening time and distance. As a result of the large populations sizes that *P. falciparum* reaches during the erythrocytic life stages we do not expect *de novo* mutations to be commonly fixed within the host, but instead expect that differences in the consensus sequences of related infections will only be fixed by passage through population bottlenecks during mosquito transmission. The lower evolutionary rate found between these two samples could be a result of long-term asymptomatic infections during which no new variants would expect to be fixed. Further investigations of the relationship between evolutionary rate and serial interval would be of great interest and may ultimately allow us to use genetic distance to infer the duration of asymptomatic infections and their contribution to local disease incidence.

Although calculations of TMRCA are difficult from such a small sample set, the placement of the basal node in the year 2000, the diversity contained within the phylogeny, and the number of persisting clades all suggest that this clonal lineage has been long-lasting and did not emerge with sample Th086.07 in 2007.

Even with this limited dataset we have successfully derived conclusions about the patterns of disease transmission in this region, but the capacity to easily derive transmission networks could be transformative for larger datasets, particularly in pre-elimination settings. Reiner *et al.* (2015) have demonstrated the capacity for network reconstruction to enable assessments of ‘malariogenic potential’ (a combined measure of disease importation rates and capacity for local transmission) in Swaziland, a country nearing elimination. The network generated by Reiner *et al.* is derived from probabilistic modeling of transmission based upon the times and geographic locations of the infections and would enable public health authorities to target interventions such as focal mass drug administration (FMDA) where they are most needed. It is worth noting that our network inference indicates that some samples will confound expectations of spatiotemporal proximity, and validation with genetic data would be a valuable addition to such mapping attempts.

In other pathogens network reconstruction using *de novo* variation has already led to actionable results for infectious-disease control and these situations could apply for malaria. There are now numerous cases in which whole-genome sequencing and network reconstruction have been deployed in real time to identify the source of a hospital-acquired infection leading directly to the identification of reservoirs of persistent disease transmission (Halachev et al. 2014; Quick et al. 2015). Even on a wider geographic scale, where comprehensive sampling of an outbreak cannot be guaranteed, the approach demonstrated here can generate concrete recommendations for public health authorities. Transmission-network reconstructions of a tuberculosis outbreak in Canada were able to show that later cases were progressions from latent to active disease, rather than active transmission, with the direct result that the outbreak was declared over, a conclusion that was reached solely by comparison of the rate of *de novo* mutation between samples (Hatherell et al. 2016). Adapting such a capacity to malaria would have major implications for elimination efforts: malaria elimination within a country is defined as zero incidence of indigenous cases (WHO 2016) and the ability to distinguish a chain of transmission from repeated importations without recourse to a multi-country dataset could be a key capacity for the WHO Global Malaria Program.

In summary, we have shown a eukaryotic parasite to be a measurably evolving population across time scales that are relevant to disease transmission, and we have calculated evolutionary rates that indicate *de novo* mutations could be found even between direct infections. This type of genetic signal is qualitatively different to those that have been available in previous genomic epidemiology studies in malaria and will enable finer grained characterization of disease transmission. Though challenges remain before these techniques can be used in higher-transmission settings, where sexual recombination will add complexity to the reconstruction, future datasets revealing *de novo* mutations in larger collections of patient samples will motivate the development of more sophisticated methods for transmission-network reconstruction. Nevertheless, there are many regions in which the techniques employed here are of immediate use. As of the end of 2015, the WHO has certified 27 countries as malaria-free, three more countries are in the process of certification (having had three years without indigenous transmission), and a further ten countries have had at least one year without indigenous transmission (WHO 2016). In any of these settings there could be outbreaks that are partially or entirely clonal. Reconstruction of a transmission network in the manner that we have achieved here would be able to detect any links that were entirely within-country, thus distinguishing imported from indigenous transmission. These data could also identify ‘super-spreader’ loci that might be targeted with focal interventions. The capacity to distinguish between indigenous and imported transmission or to pinpoint hotspots in countries nearing elimination could be key capacities for malaria elimination efforts.

## Materials and Methods

### Lab Strain Sample Preparation

DISCOVAR libraries were prepared for two Dd2 lines (Dd2-2D4 and Dd2-FDK), and one 3D7 line (3D7-Gold). Although the Dd2 lines derive from a common subcloned patient isolate, small numbers of *de novo* variants are expected to have been acquired via mutation and genetic drift in the years since the laboratory lines diverged. Three biological replicates (using separate isolates of DNA) were prepared for the DD2-FDK and 3D7-Gold samples, and one library of the Dd2-2D4. All libraries were sequenced in duplicate sequencing lanes allowing us to remove individual miscalled bases where they were not consistent across lanes.

### Senegal Sample Preparation

Samples used in this study derive from prior collections (Daniels et al. 2013; Daniels et al. 2015) in Thiès, Senegal. All samples were collected from individuals after informed consent of either the subject or a parent/guardian. This protocol was reviewed and approved by the ethical committees of the Senegal Ministry of Health (Senegal) and the Harvard T.H. Chan School of Public Health (16330, 2008) as detailed in Daniels *et al.* (Daniels et al. 2015). Samples were collected passively from patients reporting to the *‘Service de Lutte Anti-Parasitaire’* (SLAP) clinic for suspected uncomplicated malaria between approximately August and December each year. Patients with acute fevers or history of fever within the past 24 hours of visiting the clinic and with no reported history of antimalarial use were considered; they were diagnosed with malaria based on microscopic examination of thick and thin blood smears and rapid diagnostic tests (RDT), as available. All samples were collected via venous blood draw and stored as glycerolyte stocks, and an aliquot of material collected on Whatman filter paper was extracted for nucleic acid material and genotyped via 24-SNP barcode as described in Daniels *et al.* 2013. From this larger dataset, 18 samples were chosen representing single infections from clades 24, 26 and 29, (3, 5 and 10 samples respectively) based on identical genotypes observed using the 24-SNP barcode. The 18 parasite samples were culture-adapted to produce parasite DNA free of host (human) contamination. Culture times were kept to a minimum (less than 15 cycles) to preserve the integrity of the original patient material.

### Sequencing And Variant Discovery

DNA extracted from all samples was size selected to give a mean fragment length of 450 bp and sequenced using an Illumina HiSeq 2500 with a read length of 250 bp, ensuring an overlap of approximately 50 bp. Sequence was aligned to the Pf3D7_v3 reference using BWA-mem v. 0.7.12 (Li 2014). Variant discovery was then performed using DISCOVAR (release no. r52488) (Weisenfeld et al. 2014). The Pf3D7 genome was subdivided into 30 kb regions and DISCOVAR was run separately for each region. In regions where it was not possible to construct a single assembly graph for variant calling, DISCOVAR was re-run with progressively smaller regions (10, 5 and 2 Mb) of the genome. In all cases a 1 Mb overlap was allowed to control for edge effects, though variants were only included up to the inner edge of this flanking region. All libraries were sequenced and genotyped in duplicate to remove individual miscalled bases derived from sequencing error. Resulting genotypes were filtered based on Phred score (i.e. no call above 23 or any below 10 included) and hypervariable sites were removed.

For comparison of this variant discovery method with other approaches, the 250 bp reads were clipped to 100 bp and aligned using the same version of BWA-mem (Li 2014 REF). Variant discovery was performed using GATK HaplotypeCaller followed by quality score recalibration using GATK VQSR (DePristo et al. 2011). The VQSR truth-set consisted of variant loci that had been previously determined in the *‘Pf-Crosses’* dataset generated by Miles *et al.* (Miles et al. 2016). This dataset consists of a series of genetic crosses of *P. falciparum* clones (3D7 x HB3, HB3 x Dd2 and 7G8 x GB4) in which parents and all offspring were deeply sequenced, enabling variant calls to be filtered according to conformity with Mendelian inheritance expectations.

### Variant Validation

The *P. falciparum* genome presents a challenge to many genotype callers due to extensive repeat structure and recombination within some large gene families (Bopp et al. 2013), as well as the high A/T nucleotide composition of the intergenic regions (Gardner et al. 2002). Moreover, validation of variant calls in low-complexity regions of the *P. falciparum* genome via conventional PCR and Sanger dideoxy sequencing is difficult, as PCR amplicons evaluated were generally too repetitive to yield interpretable sequence traces. As a result, a multi-stage concordance-based approach to genotyping was taken: 1) the ‘accessible’ genome was determined separately for each genotype caller; 2) genotypes in the regions accessible to both callers were compared to a known sample; 3) genotypes in difficult to access regions were compared to those in accessible regions.

#### Accessible Genome Determination

Miles *et al.* used the *Pf-Crosses* dataset in order to characterize the ‘accessible genome’ – the region in which genotyping calls could be reliably made (Miles et al. 2016). The authors found 90% of the genome to be within an acceptable error rate, and only hypervariable regions, centromeres and telomeres were excluded. DISCOVAR generates few variables upon which variant recalibration could be performed; therefore, we took a stricter approach. Using *in silico* sequence generated from a second genome assembly, we determine a region to be accessible only if the variant caller can successfully reconstruct the region around all variants within it.

An artificial *(in silico)* DISCOVAR-appropriate Dd2-2D4 read set (250 bp fragment size, 450 bp insert size) was generated from a PacBio assembly of the Dd2-2D4 line of *P. falciparum* (PfDd2_v1) using wgsim v0.3.2 (Li et al. 2009). This read set was aligned to the standard Pf3D7_v3 reference assembly and genotypes were called using DISCOVAR as detailed above. For each resulting variant, the putative Dd2-2D4 sequence in a 200 bp region around the variant was reconstructed and realigned to the Dd2-2D4 PacBio reference via Smith-Waterman alignment (BWA-SW (Li and Durbin 2010)). Any differences between this locally reconstructed Dd2-2D4 sequence and the Dd2-2D4 PacBio assembly (i.e. the Levenshtein distance between the two strings) would therefore represent errors derived solely from the DISCOVAR algorithm. Calls were also made using an artificial GATK HaplotypeCaller (DePristo et al. 2011) dataset (100bp reads, 300bp insert size) and assessed using the same Smith-Waterman approach. For each variant calling method, the ‘accessible genome’ was defined as those regions where alignment depth was within one standard deviation of the mean and where the Levenshtein distance (LD) was zero (i.e., where reconstruction of Dd2-2D4 had been error free).

#### Accessible Marker Validation

An experimentally validated set of genotypes was then used for confirmation of variants within those regions of the genome that were accessible to both HaplotypeCaller and DISCOVAR. This biological validation was performed using three laboratory lines of *P. falciparum,* each sourced from different laboratories. The called variants were then compared to the set of previously validated *‘Pf-Crosses’* variants using VCFEval (Cleary et al. 2015): receiver-operator curves (ROC curves – i.e. plots of true positive and false positive rates at equal quality scores) and specificity/sensitivity values were calculated in the regions that had been previously determined to be accessible by both genotype callers and in the regions determined to be accessible only to DISCOVAR.

#### ‘Inaccessible’ Marker Validation

In regions of the genome determined to be accessible only to DISCOVAR, pairwise genetic distances between the Senegal samples were compared from “DISCOVAR-only” regions to those regions in the core genome that were callable with both methods. Although individual validation of markers is difficult due to a lack of suitable examples within these regions, we expect correlation between pairwise distances, such that pairs with closely related accessible genomes should also have closely related inaccessible genomes, and distantly related accessible genomes will have distantly related inaccessible genomes.

### Phylogeny & Transmission Networks

SNP and INDEL distance matrices were calculated based on the number of mutations between each sample pair, without making adjustments based on putative mutation rate or STR length. Parsimony trees were generated using the R package *Phangorn* (Schliep 2011) and neighbor-joining trees were generated using the R package ‘ape’ (Didier et al. 2015). Phylogenies were calculated by both methods for the pairwise SNP distance, INDEL distance and the Manhattan distance of both SNPs and INDELs. Bootstrapping was performed by ‘sampling with replacement’ for 100 replicates across each phylogeny. Bootstrap support is reported as the proportion of trees supporting each node.

To assess the phylogenetic signal within the clades the Tree-Puzzle program (Schmidt et al. 2002) was used to analyze all potential sequence quartets within the set. For any set of four individuals, a maximum of three phylogenies are possible, and the TreePuzzle method randomly selects individuals from the dataset and assesses by maximum likelihood if it accords with any of the three phylogenies (see Strimmer *et al.* for details (Strimmer and von Haeseler 1996)). A quartet with clean phylogenetic signal will show high likelihood to one of the tree topologies, a network with significant introgression will show likelihood to more than one, and an unresolved star-like phylogeny will show likelihood to none.

Temporal data were assessed by examining the correlation between sampling time and the root-to-tip distance of a ‘non-clock’ phylogenetic tree (in this case one derived by neighbor-joining) using TempEST v1.5 (Rambaut et al. 2016). Correlation between genetic distance and time was assessed via Pearson’s product moment. Transmission networks were reconstructed using SeqTrack within adegenet (Jombart et al. 2011) based on the Manhattan distance of all variants an approach that has been shown to outperform parsimony based methods where sampling density is low (Worby et al. 2017). Bootstrapping of the transmission network was performed by sampling all variants with replacement for 100 iterations; bootstrap values are reported as the proportion of bootstrapped networks supporting each join.

### Evolutionary Rate Calculation

For samples sharing very recent common ancestry, pairwise variant call differences are expected to be dominated by sequencing error, leading to overestimation of the rate at which *de novo* mutations accumulate. Where a transmission network could be resolved for a clade, evolutionary rates were therefore calculated for each IBD pair as the difference between the distance of each sample to an outgroup (chosen from one of the other two clades) to account for ancestral diversity (Li et al. 1988). This was performed separately for SNPs and INDELs yielding a separate evolutionary rate for each variant type, and again within the regions of the genome determined to be callable with both callers and the DISCOVAR-only genome yielding an estimate of the contribution to the evolutionary rate of ‘accessible’ and ‘inaccessible’ regions of the genome. The mean evolutionary rate is reported for each pair of samples using all potential outgroups.

## Acknowledgements

We are grateful for the assistance of Thomas D Otto (Sanger Institute) for providing the DD2_v1 reference genome for validation. David Fidock (Columbia University) and Daniel Goldberg (Washington University in St Louis) for supplying their *Plasmodium* stocks for sequencing. We thank the UCAD/LeDantec laboratory and SLAP clinic team, particularly Dr Ngayo Sy, and the patients for clinical samples. We thank Sinéad Chapman, Caroline Cusick, and Jim Bochicchio for project management, and Steve Schaffner and Angela Early for their insightful comments on the manuscript. Funding for this work at the Harvard TH Chan School of Public Health and the Broad Institute was provided by a grant to DF Wirth from the Global Health Program at the Bill and Melinda Gates Foundation.

